# Experimental Suppression of TMS-EEG Sensory Potentials

**DOI:** 10.1101/2022.02.02.478881

**Authors:** Jessica M. Ross, Manjima Sarkar, Corey J. Keller

## Abstract

**Background:** The sensory experience of transcranial magnetic stimulation (TMS) evokes cortical responses measured in EEG that confound interpretation of TMS-evoked potentials (TEPs). Methods for sensory masking have been proposed to minimize sensory contributions to the TEP, but the most effective combination for suprathreshold TMS to dorsolateral prefrontal cortex (dlPFC) is unknown.

**Objective:** We applied sensory suppression techniques and quantified electrophysiology and perception from suprathreshold dlPFC TMS to identify the best combination to minimize the sensory TEP.

**Methods:** In 21 healthy adults, we applied single pulse TMS at 120% resting motor threshold (rMT) to the left dlPFC and compared EEG vertex N100-P200 and perception. Conditions included three protocols: *No masking* (no auditory masking, no foam, jittered inter-stimulus interval (ISI)), *Standard masking* (auditory noise, foam, jittered ISI), and our ATTENUATE protocol (auditory noise, foam, over-the-ear protection, unjittered ISI).

**Results:** ATTENUATE reduced vertex N100-P200 by 56%, “click” loudness perception by 50%, and scalp sensation by 36%. We show that sensory prediction, induced with predictable ISI, has a suppressive effect on vertex N100-P200, and that combining standard suppression protocols with sensory prediction provides the best N100-P200 suppression. ATTENUATE was more effective than *Standard masking*, which only reduced vertex N100-P200 by 22%, loudness by 27%, and scalp sensation by 24%.

**Conclusions:** We introduce a sensory suppression protocol superior to *Standard masking* and demonstrate that using an unjittered ISI can contribute to minimizing sensory confounds. ATTENUATE provides superior sensory suppression to increase TEP signal-to-noise and contributes to a growing understanding of TMS-EEG sensory neuroscience.

**Highlights:**

- ATTENUATE is a novel sensory suppression protocol for suprathreshold dlPFC TMS
- ATTENUATE is superior to standard masking for minimizing sensory confounds
- ATTENUATE reduced vertex N100-P200 by 56% with no effect on the early TEP
- ATTENUATE reduced “click” loudness rating by 50% and scalp sensation by 36%
- Individual modifications are not sufficient to reduce vertex N100-P200 or perception

## 1. Introduction

Transcranial magnetic stimulation (TMS) is a powerful non-invasive tool for stimulating brain networks [1–3] and has proven useful for the neurophysiological characterization and treatment of neurological and psychiatric disorders [4–8]. Neural changes caused by TMS are measurable and quantifiable using electroencephalography (EEG) [9–13]. For instance, an averaged single pulse TMS-evoked EEG potential (TEP) can be used to characterize local and network excitability as well as plasticity following repetitive TMS protocols [10,14–16]. Gaining a better understanding of and utilizing TMS-induced EEG changes is critical for targeted and personalized circuit manipulation for robust clinical use.

While TEPs are a promising measure of TMS-evoked neural activity, it has become evident that off-target sensory effects of single TMS pulses can severely confound the interpretation of the TEP [17–22]. These off-target effects include sensory potentials that are peripherally evoked due to the multisensory nature of TMS [23]. Although the TEP is reproducible [9,24] and has been shown to reflect localized TMS-evoked activity at the earliest latencies after the pulse (up to approximately 60-80ms) [11,16,20,23,25–27], there is accumulating evidence that the later TEP (>80ms) is contaminated by off-target sensory potentials [18–20]. One such component of the later TEP is an evoked response [28] induced from the sound of TMS (referred to as the auditory evoked potential, AEP) and not specific to the site of stimulation [17,23]. The greatest amplitude and most robustly measured subcomponents of this sensory potential occur at the vertex at ∼100 and 200 ms with an accompanying smaller potential at ∼50 ms [22,25,29–33]. These vertex potentials are described as an N100-P200 complex, which overlaps with all but the earliest TEP components. In summary, sensory potentials in the TEP remain a significant confound to the direct effects of TMS and minimization or removal is necessary to improve interpretability.

Experimental modifications have been proposed to suppress the sensory vertex N100-P200, but the most effective combination for suprathreshold TMS is unknown. This is particularly true for targeting the dlPFC, the primary treatment location for many neuropsychiatric disorders [34–39]. Here, we focus on the following experimental modifications: auditory masking, a foam separator between the coil and the scalp, and predictably spacing TMS pulses. A common sensory masking protocol is to pair earplugs and/or auditory noise masking [2,19], a foam separator, and a jittered inter-stimulus interval (ISI) (hereafter called *Standard masking*). Recent evidence for effective standard masking is promising for subthreshold TMS to primary motor cortex (90% of the resting motor threshold (rMT) [2,19]). However, it has also been shown that these methods often do not fully suppress the sensory vertex N100-P200 [17,18,40–44], particularly for higher intensity protocols [17,40]. Rocchi *et al*. [19] used over-the-ear protection in addition to noise masking to further minimize vertex N100-P200, with positive results for subthreshold M1 stimulation. However, how over-the-ear protection performs for higher stimulation intensities and non-M1 targets is unknown. The use of foam padding between the coil and scalp is thought to suppress the vertex N100-P200 by reducing bone conduction of the sound [41]. However, it is unclear what type or thickness of foam should be used. In addition, there is no consensus regarding how to adjust stimulation intensity to account for higher coil to cortex distance when foam is added. Modifying the inter-stimulus interval (ISI) timing changes the predictability of TMS pulses, which can have an effect on MEP amplitude [45,46]. However, whether more predictable TMS timing results in a similar attenuation of the TEP is unknown. In summary, a thorough investigation into the optimal experimental methodology to suppress sensory vertex N100-P200 following suprathreshold TMS to the clinically significant dlPFC is necessary.

In this study, we develop an optimal combination of experimental modifications that maximally reduce the vertex N100-P200 complex and sensory perception following suprathreshold single pulse TMS to the dlPFC. In a sample of 21 typically healthy adults, we compared the effects of three masking protocols –: *No masking, Standard masking*, and a novel procedure – on the vertex N100-P200 and on perception of the TMS loudness, scalp sensation, and pain. We hypothesized that our novel combination of experimental procedures, with the addition of further sound dampening and modification of TMS timing, would best suppress the non-specific sensory component of the TEP. This work contributes to a growing understanding of TMS-EEG sensory neuroscience, and the novel protocol has the potential to enhance interpretability of TMS-EEG studies.

## 2. Methods

### 2.1. Participants and Study Design

All data were collected at Stanford University under an approved institutional review board protocol after participants gave their written informed consent. Participants (N=21) were 19-64 years old (44.0 mean +/- 14.58 SD) and without current psychiatric or neurological diagnoses. A wide age range was chosen so as not to constrain findings to any a priori group. Supplementary table S1 includes demographic information for all subjects. For each participant, the experiment was conducted on a single day. The experiment was split into multiple single pulse TMS-EEG blocks. Each block consisted of 80 individual single pulse TMS trials applied to the left dlPFC. 80 trials were chosen as they provided high test-retest reliability of the N100 and P200 [9]. TMS-evoked potentials (TEPs) and perceptual scores were quantified, as described below and in schematic in Figure 1A.

**Figure 1.**
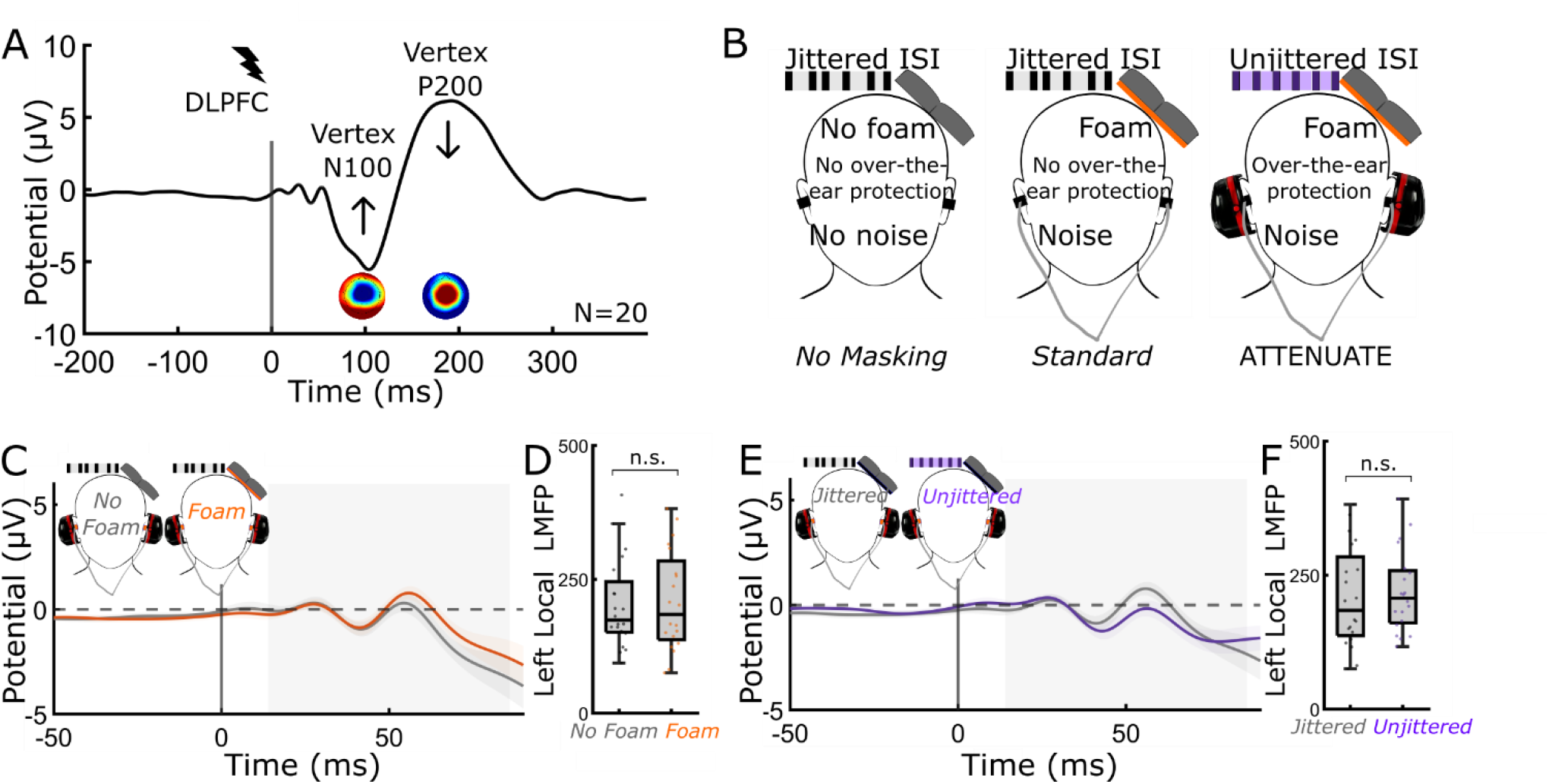
Experimental design. (A) Single pulse TMS-EEG to the left dlPFC. TMS was applied at 120% rMT (80 trials). Perceptual reports of loudness of the TMS “click”, intensity of scalp sensation, and pain followed each condition. Vertex N100-P200 was quantified using LMFP. Arrows denote that the goal of the study was to minimize the N100-P200 and sensory perception while preserving the early TEP. (B) Three experimental conditions were compared: *No masking* (jittered ISI/no foam/no noise/no over-the-ear protection), *Standard masking* (jittered ISI/foam/noise/no over-the-ear protection), and ATTENUATE (unjittered ISI/foam/noise/over-the-ear protection). (C-F) Neither *foam* (C-D) nor *unjittered ISI* (E-F) altered the early local TEP (14-86 ms). (C-D) Effect of *foam* on early (14-86ms) local TEP. *Foam* did not modify the early local TEP (T = -0.35, DF = 19, p = 0.73). (E-F) Effect of using an *unjittered ISI* on early local TEP. Modifying the timing of TMS pulses did not change the early local TEP (T = -0.58, DF = 19, p = 0.57). All error bars denote standard error. N.S. = not significant.

### 2.2. Transcranial Magnetic Stimulation

TMS was performed with a MagPro X100 stimulator (MagVenture, Denmark) and a MagVenture Cool B65 figure-of-eight coil (MagVenture, Denmark). The motor hotspot for the right first dorsal interosseous (FDI) was determined by delivering single TMS pulses to the left motor cortex. Resting motor threshold (rMT) was obtained once at the beginning of the experiment and defined as the intensity that produced a visible twitch in relaxed FDI in ≥ 5/10 stimulations [47,48]. Neuronavigation (Localite TMS Navigator MR-less system, Alpharetta, GA) was utilized to determine the left dorsolateral prefrontal cortex (dlPFC) location on a standard Montreal Neurological Institute (MNI) brain map, fitted to individual participants’ heads based on scalp measurements. The left dlPFC site (MNI -38, 22, 38) was used to target the fronto-parietal control network [49]. TMS coil angle was placed at the angle between 0 and 90 degrees [50–54] that most minimized discomfort and pain for each subject (M=52 degrees, SD=27). Supplementary table S1 includes the optimal angle for each subject.

To identify the set of procedures that maximally reduces the vertex N100-P200, we tested our novel combination of experimental procedures, which we refer to as <u>ATTENUATE</u> (<u>A</u>uditory: noise masking, <u>T</u>iming: unjittered ISI, <u>T</u>actile: foam, and over-the-<u>E</u>ar protection to <u>N</u>egate <u>U</u>nwanted <u>A</u>rtifacts in <u>T</u>MS-<u>E</u>EG). We tested ATTENUATE against a common *Standard masking* procedure (auditory noise, foam, jittered ISI) and *No masking* (no auditory noise, no foam, jittered ISI). See Figure 1B for schematics of the three masking procedures. As noted in the *Introduction*, it is becoming increasingly common in TMS-EEG studies to employ the *Standard masking* protocol that uses foam and a “click” frequency auditory masking noise [17,19].

In follow-up contrasts we quantify the effectiveness of auditory masking (*auditory noise* with and without *over-the-ear protection*), foam, and ISI timing modifications alone (i.e. when each of the other factors is held constant). Figure 3A,H,O depict auditory, foam, and ISI timing conditions, respectively. Conditions were presented in a pseudorandomized order. All conditions were collected in each subject unless the experiment ended early due to time constraints. Supplementary tables S2-S3 reflect for each subject the conditions performed.

Each TMS-EEG condition consisted of 80 single pulses (biphasic pulses at 280μs pulse width) at an intensity of 120% rMT. Stimulator recharge was delayed to 500ms to prevent recharge artifact from affecting EEG in the time period of interest [55]. Participants were instructed to keep their eyes open and gaze relaxed throughout each run. For conditions using auditory noise, the noise sound matched the frequency of the TMS click [11] and was delivered with earplug earbuds (Elgin USA Ruckus Earplug Earbuds, NRR 25 dB, Arlington, Texas) at the maximum volume comfortable for each participant. In conditions using *over-the-ear protection*, to further dampen the TMS “click” sound before reaching the ear canal, we used over-the-ear noise-reducing foam-filled earmuffs (3M Ear Peltor Optime 105 behind-the-head earmuffs, NRR 29 dB, Maplewood, Minnesota). In conditions without auditory noise, earplug earbuds were still kept in the ear canals but no noise was played. In conditions requiring foam, a thin (0.5 cm) foam pad was attached to the TMS coil, and rMT was redetermined using this foam to accurately deliver a TMS intensity at 120% rMT while accounting for the increase in coil to scalp distance (see [23] for effects of separator and of distance to scalp on amplitude of vertex N100-P200). Supplementary table S1 includes all rMTs, with and without foam, for all subjects. To determine whether predictability of TMS pulses can attenuate sensory components in the TEP, we compared jittered ISI (2±1 s jitter) and unjittered ISI (2 s) protocols.

### 2.3. Electroencephalography

64-channel EEG data were obtained using a BrainVision actiCHamp Plus amplifiers (5 kHz sampling rate), with ActiCAP slim active electrodes in an extended 10-20 system montage (actiCHamp, Brain Products GmbH, Munich, Germany). EEG data were online referenced to Afz, recorded using BrainVision Recorder software v1.24.0001 (Brain Products GmbH, Germany). Impedances were maintained below 5 kΩ.

#### 2.3.1. Preprocessing of TEPs

All EEG preprocessing and analyses were performed in MATLAB R2021a (Mathworks, Natick, MA, USA) using the EEGLAB v2021.1 toolbox [56] and custom scripts. Removal of artifactual EEG data was performed using a custom preprocessing pipeline, as is most common [57], but followed most closely with Ross *et al*. [40] (steps prior to sensory removal), TMSEEG [58], and TESA [59]. Due to a marked impact of preprocessing pipelines on the TEP, as reported in [57], we took a conservative approach in all steps that required human judgement (with minimal data deletion) and describe all preprocessing steps used in detail with justification for each choice and supporting literature.

All details of EEG data cleaning can be found in Supplementary section S1.1.

#### 2.3.2. Quantification of TEPs

For time window and region of interest (ROI) selection, and calculation of global mean field power (GMFP) and local mean field power (LMFP), see Supplementary section S1.2 and Figures S2-5. To compare vertex N100-P200 across experimental conditions, TEPs were generated as averages over the vertex ROI: FC1, FCz, FC2, C1, Cz, C2 (Figure S5C for ROI). LMFP was calculated for the ROI and the area under the curve (AUC) of the LMFP was quantified for the appropriate time windows. Supporting that our time windows and ROI capture the vertex N100-P200 complex, we observed a strong correlation between vertex N100 and P200 (Figure S7A,B; area under the curve (AUC) of LMFP; r(19)=0.91, p=0.00000004; regression: F(1,19)=81.68, p=0.00000004; R2 = 0.82).

To verify that sensory suppression techniques did not alter the early local TEP, we compared LMFP of the early window (14-86 ms) in electrodes local to the site of stimulation. For each condition, the AUC of the LMFP was utilized (and referred to simply as LMFP in the manuscript). Contrasts included *No foam* vs. *Foam* conditions (with other factors matched) and *Jittered* vs. *Unjittered* conditions (with other factors matched). To identify an ROI for examining local response to TMS, electrodes maximally different from baseline in the early window were chosen: AF3, AFz, F3, F1, FC3, FC1 (Section S1.2 and Figure S6C for ROI). We found no significant effect on the early TEP response (LMFP) of using *Foam* (T = -0.3534, DF = 19, p = 0.7277) or an *Unjittered* protocol (T = -0.5773, DF = 19, p = 0.5705; Figure 1C-F).

To ensure that an unjittered ISI did not induce changes in the early local TEP over time, we compared the first half of trials to the second half in the *Unjittered* condition [60,61]. We observed no significant difference in the early LMFP between the first half and second half of trials in the *Unjittered* (2 second ISI) condition (Figure S6; T=1.2542, DF=19, p=0.2250, CI=-11.4509, 45.6910).

#### 2.3.3. Statistical analyses of TEPs

To compare single pulse TEP responses across the three masking protocols, we computed the LMFP for the N100 and P200 time windows in the central ROI. We performed an Analysis of Variance (ANOVA, repeated measures) with three levels (*No masking, Standard masking*, ATTENUATE), followed by post hoc pairwise comparisons using Tukey’s HSD procedure where appropriate.

### 2.4. Perceptual Ratings

To assess perceptual experience during each stimulation condition, participants were asked to respond verbally immediately following each condition to rate *loudness, scalp sensation*, and *pain perception* on scales ranging from 0 to 10. These scores were inputted into the research electronic data capture system (REDCAP, Vanderbilt, Nashville, TN). To ensure consistency in how these questions were phrased across conditions and subjects, the following scripts were used:

*With 0 being you could not hear it, and 10 being as loud as a fire alarm, how loud did you perceive the ‘click’ sound to be?*

*With 0 being you could not feel it, and 10 being it felt as intense as a hard flick, how much did you feel the tapping sensation?*

*With 0 being no pain at all, and 10 being unbearable pain, how much pain did you feel?*

#### 2.4.1. Statistical analyses of perceptual ratings

Raw perceptual ratings were compared across the three conditions using a repeated measures ANOVA with three levels (*No masking, Standard masking*, ATTENUATE), followed by post hoc pairwise comparisons using Tukey’s HSD procedure where appropriate.

### 2.5. Interactions between perceptual ratings and vertex N100-P200

To further understand the relationship between perceptual ratings of loudness, scalp sensation, pain, and the vertex N100-P200, an exploratory analysis compared perceptual ratings and vertex N100-P200 LMFP values across subjects for the *No masking* condition only. The goal of this analysis was to better understand the relationship between unsuppressed sensory contributions. For this analysis, a Pearson’s correlation matrix was generated with correlation coefficients (Figure S7), and follow-up linear univariate regression analyses were performed for significantly correlated factors using the *regress* function in MATLAB R2021a (Mathworks, Natick, MA, USA) [62,63].

## 3. Results

### 3.1. The ATTENUATE protocol is superior to *Standard masking* at reducing the vertex N100-P200

The vertex N100 and P200 LMFPs (see Section 2.3.2 above for ROI and window selection) were compared across the three sensory suppression protocols (*No masking, Standard masking*, and ATTENUATE). Vertex LMFP was significantly different across conditions in the N100 (F(2,57)=3.64, p=0.03) and P200 (F(2,57)=9.40, p=0.0003) time windows (Figure 2B-D). Post-hoc pairwise comparisons revealed that ATTENUATE reduced the LMFP vertex N100 (M=145.84, SD=62.13) compared to *No masking* (M=304.28, SD=288.54; Tukey’s HSD, p=0.03). *Standard masking* did not show statistical differences from *No masking* (M=236.38, SD=167.59; p=0.52) or from ATTENUATE (p=0.26). ATTENUATE reduced the LMFP vertex P200 (M=311.98, SD=130.51) compared to *No masking* (M=727.68, SD=401.13; p=0.0002) and compared to *Standard masking* (M=563.90, SD=309.63; p=0.03). *Standard masking* did not show statistical differences from *No masking* (p=0.21). These results reflect that ATTENUATE reduced vertex N100 by 54.41% and vertex P200 by 56.58% from *No masking* (average of 55.94% reduction across the vertex N100-P200 complex). In comparison, *Standard masking* reduced vertex N100 by 22.31% and vertex P200 by 22.51% (average of 22.45% across the vertex N100-P200 complex). In summary, we observed a significant group effect across sensory suppression procedures and ATTENUATE reduced the vertex N100 and P200 more than *Standard masking*.

**Figure 2.**
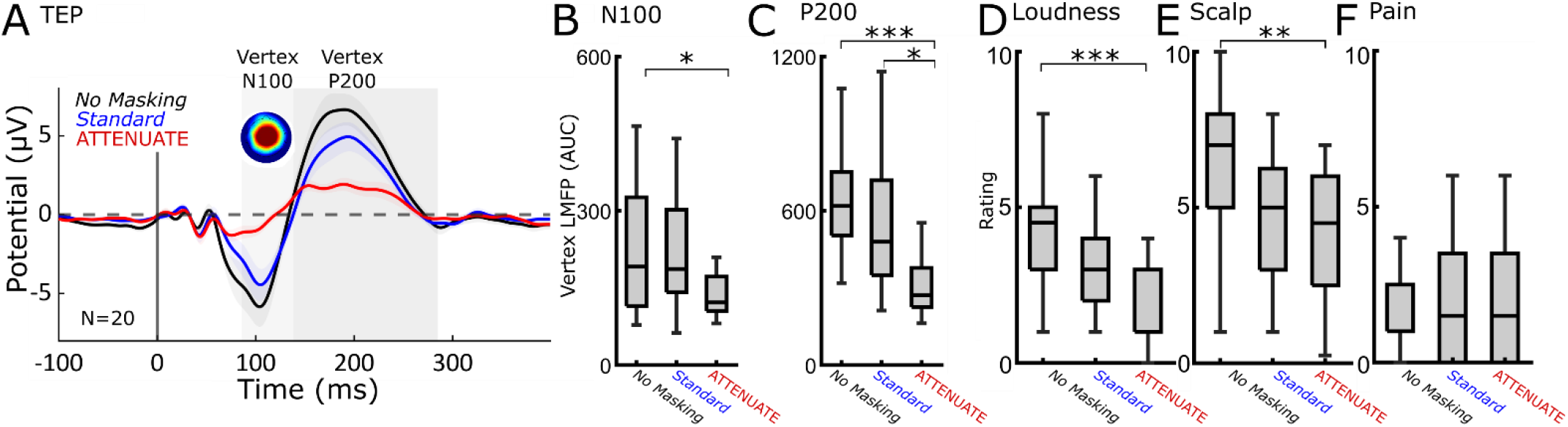
The ATTENUATE protocol is superior to *Standard masking* at reducing vertex N100-P200, “click” loudness perception, and scalp sensation. (A) Group mean TEPs of vertex ROI (N=20). Shaded areas indicate time windows used for analysis. (B-C) ATTENUATE reduces the vertex N100-P200. Vertex LMFP is reduced in both N100 (F(2,57)=3.64, p=0.03) and P200 (F(2,57)=9.40, p=0.0003) time windows across the three conditions. Pairwise comparisons revealed that ATTENUATE reduced vertex N100-P200 compared with *No Masking* (N100: p=0.03; P200: p=0.0002), and that ATTENUATE reduced vertex P200 compared with *Standard masking* (p=0.03). Comparisons between *No Masking* and *Standard masking* were non-significant (N100 p=0.52; P200 p=0.21). (D-E) Sensory suppression protocols reduced perception of loudness (F(2,57)=8.53, p=0.0006) and scalp sensation (F(2,57)=5.47, p=0.007). Pairwise comparisons demonstrate that ATTENUATE reduced both loudness (loudness p=0.0004) and scalp sensation (scalp sensation p=0.006) from *No masking*, but did not reduce compared with *Standard masking* in loudness rating (p=0.15) or scalp sensation (p=0.56). Comparisons between *No masking* and *Standard masking* were non-significant (loudness p=0.07, scalp sensation p=0.08). (F) Pain was not effected by protocol (F(2,57) = 0.06, p = 0.9461). All significant pairwise comparisons are indicated with brackets and asterisks mark level of significance. *p<.05, ** p<.01, *** p<.001. All error bars denote standard error.

### 3.2. ATTENUATE is more effective than *Standard masking* at reducing loudness perception and scalp sensation

Raw perceptual ratings for loudness of “click”, sensation on the scalp, and pain were compared across the three masking protocols (*No masking, Standard masking*, and ATTENUATE). See Figure 2A for perceptual ratings following *Standard masking* and ATTENUATE conditions. We found a significant difference in loudness perception (F(2,57)=8.53, p=0.0006) and scalp sensation (F(2,57)=5.47, p=0.0067) across conditions (Figure 2A). Perception of pain did not change between the conditions (F(2,57) = 0.06, p = 0.9461). Post-hoc pairwise comparisons revealed that ATTENUATE reduced loudness rating (M=2.24, SD=1.51; p=0.0004) and scalp sensation (M=4.15, SD=2.10; p=0.006) compared to *No masking* (Loudness: M=4.23, SD=1.92; Scalp: M=6.30, SD=2.36). *Standard masking* (Loudness: M=3.08, SD=1.51; Scalp: M=4.78, SD=2.07) did not show a statistical difference from *No masking* (Loudness: p=0.07; Scalp: p=0.08) or from ATTENUATE (Loudness: p=0.15; Scalp: p=0.56). These results reflect that ATTENUATE reduced loudness rating by 50.30% and scalp sensation by 35.52% from *No masking*. In comparison, *Standard masking* reduced loudness rating by 27.22% and scalp sensation by 24.21%. In summary, we observed a significant group effect across sensory suppression procedures with ATTENUATE reducing the perception of “click” loudness and scalp sensation compared with *No masking*.

### 3.3. Individual auditory, foam, or ISI timing modifications are not sufficient for reducing vertex N100-P200 or sensory perception

To determine if components of these sensory suppression modifications in isolation reduce the vertex N100-P200 or sensory perception, we compared vertex N100-P200 LMFP and perceptual ratings across auditory (*No noise, Noise, Noise and over-the-ear protection*; Figure 3A-D), foam (*No foam, Foam*; Figure 3E-H), and ISI timing (*Jittered, Unjittered*; Figure 3I-L) conditions. For each comparison, all other modifications were matched.

**Figure 3.**
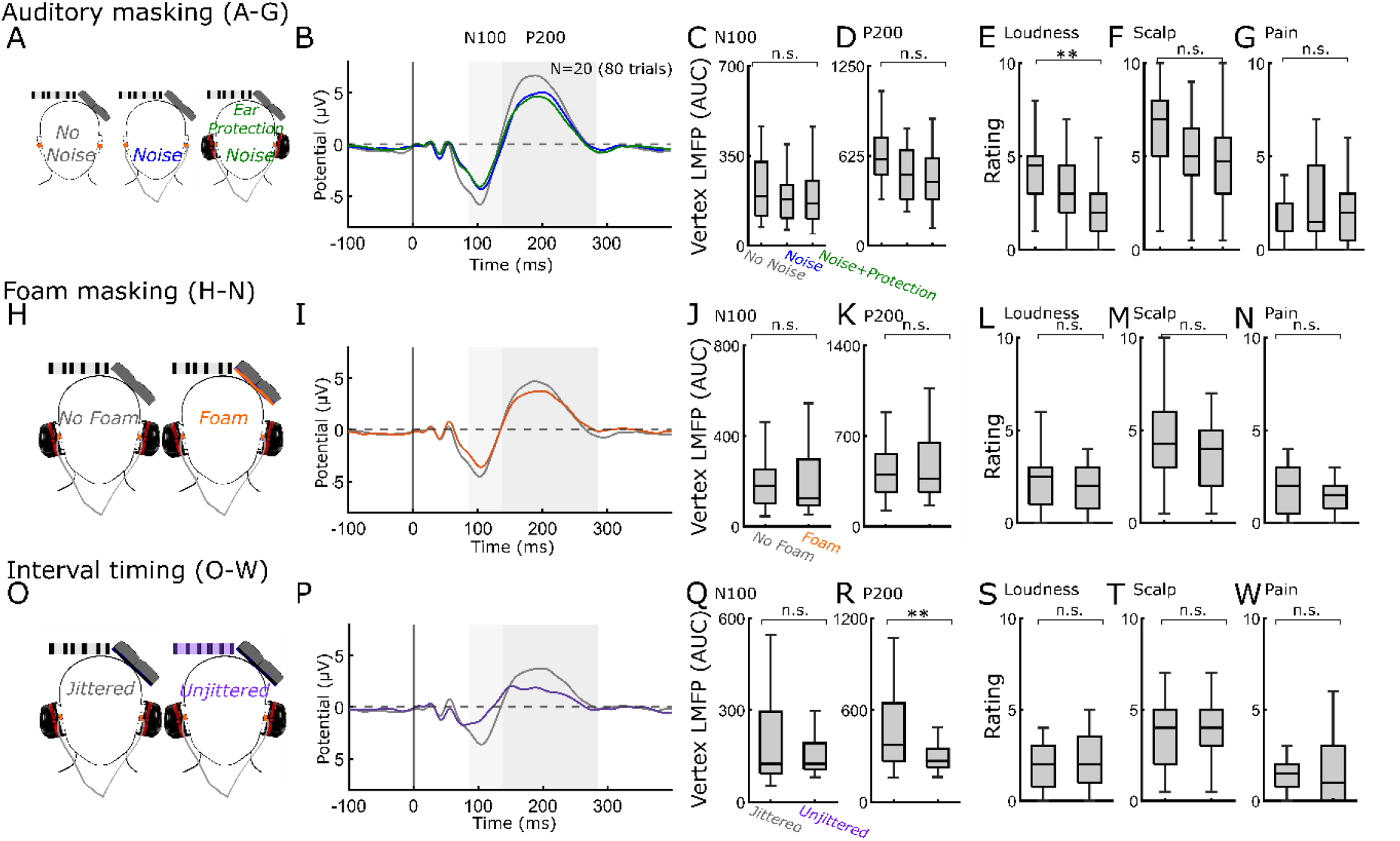
Individual auditory, foam, or timing modifications are only minimally effective for reducing vertex N100-P200 or sensory perception. (A) Auditory masking conditions: *No noise, Noise, Noise with over-the-ear protection* (B) TEPs from the vertex ROI (C-D) LMFP for N100 (C) and P200 (D) time windows. ANOVAs revealed that auditory masking protocols only had a marginal but insignificant effect on vertex P200. (E-G) Perceptual ratings of “click” loudness (E), scalp sensation (F), and pain (G). ANOVAs revealed that auditory masking protocols had an effect on loudness rating, driven by *Noise with over-the-ear protection* change from *No masking*, an insignificant reduction in scalp sensation, and no effect on pain. (H) *No foam* and *Foam* conditions (I) TEPs from the vertex ROI (J-K) LMFP. T-tests revealed that *Foam* had no effect on vertex N100 or P200. (L-N) Perceptual ratings of loudness (L), scalp sensation (M), and pain (N). T-tests revealed that *Foam* had no effect on any perceptual ratings. (O) *Jittered* and *Unjittered* ISI conditions (P) TEPs from the vertex ROI (Q-R) LMFP. T-tests revealed that using an *Unjittered ISI* had a non-significant effect in vertex N100, and a significant effect on vertex P200. (S-W) Perceptual ratings of loudness (S), scalp sensation (T), and pain (W). T-tests revealed that using an *Unjittered* ISI had no effect on any perceptual ratings. p<.05, ** p<.01, *** p<.001. All error bars denote standard error. Shaded areas indicate time windows used for analysis.

#### 3.3.1. Auditory suppression

An ANOVA across the three auditory conditions (*No noise, Noise, Noise and over-the-ear protection*) revealed no effect on vertex N100 (Figure 3C; F(2,57)=1.27, p=0.29) with an insignificant but marginal effect on vertex P200 (Figure 3D; F(2,57)=3.10, p=0.05). Auditory suppression had an effect on loudness rating across the three conditions (Figure 3E; F(2,57)=6.12, p=0.0039). Post-hoc pairwise comparisons revealed that *Noise with over-the-ear protection* reduced loudness (M=2.25, SD=1.59) compared to *No noise* (M=4.23, SD=1.92; Tukey’s HSD; p=0.003). *Noise* alone did not reduce loudness rating (M=3.10, SD=1.84) from *No noise* (p=0.12) but was also not different from *Noise with over-the-ear protection* (p=0.30). Auditory suppression protocols did not reduce scalp sensation (Figure 3F; F(2,57)=3.08, p=0.05) or pain rating (Figure 3G; F(2,57) = 0.56, p = 0.5730).

#### 3.3.2. Foam

The use of *Foam* had no effect on vertex N100 (Figure 3J; T = -0.2886, DF=19, p = 0.7760, CI= -46.9654, 35.5838) or vertex P200 (Fig 3K; T = -0.8277, DF=19, p = 0.4181, CI= - 122.5488, 53.0926). *Foam* also had no effect on loudness rating (Figure 3L; T = 1.0918, DF=19, p = 0.2886; CI= -0.3210, 1.0210), scalp sensation (Figure 3M; T = 1.4690, DF=19, p = 0.1582; CI= -0.3398, 1.9398), or pain rating (Figure 3N; T = 1.5305, DF=19, p = 0.1424; CI= -0.2114, 1.3614).

#### 3.3.3. ISI Timing

Using an *Unjittered ISI* had a non-significant suppressive effect on vertex N100 (Figure 3Q; T = 1.8574, DF=19, p = 0.0788, CI= -7.5349, 126.3170) and a significant suppressive effect on vertex P200 (Figure 3R; T = 3.8362, DF=19, p = 0.0011, CI= 78.0444, 265.4598). *Unjittered ISI* had no effect on loudness rating (Figure 3S; T = -0.8193, DF=19, p = 0.4228; CI= -0.8443, 0.3693), scalp sensation (Figure 3T; T = -1.1981, DF=19, p = 0.2456; CI= - 1.6482, 0.4482), or pain rating (Figure 3W; T = -0.0901, DF=19, p = 0.9291; CI= -0.6056, 0.5556).

In summary, *auditory, foam*, or *ISI timing* modifications alone are only minimally effective strategies for reducing vertex N100-P200 LMFP and perceptual ratings of “click” loudness, scalp sensation, or pain.

### 3.4. Pain may be contributing to vertex N100-P200

Finally, to better understand the relationship between electrophysiology and perception, we compared the vertex N100, vertex P200, and perceptual ratings in the *No masking* condition (Figure S7) using a correlation matrix of all measures. All relationships were insignificant except for between N100 and P200 as well as between Pain and the P200 (Figure S7A; Methods section 2.3.2). Pain rating had a positive correlation with vertex P200 (Figure S7C; correlation: r(19)=0.45, p=0.04; regression: F(1,19)=4.67, p=0.04; R^2^ = 0.20).

## 4. Discussion

In the present study, to reduce the sensory effects of TMS, we sought to experimentally minimize the vertex N100-P200 and sensory perception arising from suprathreshold TMS to the dlFPC. We developed a novel combination of experimental sensory suppression techniques, termed ATTENUATE, which consisted of auditory noise masking, foam, over-the-ear protection, and unjittered pulse timing. To our knowledge, this is the first study to present the ATTENUATE protocol. We find the following: 1) The ATTENUATE protocol significantly reduced the vertex N100-P200 by 56%, outperforming other standard masking procedures, with no effect on the early TEP; 2) The ATTENUATE protocol reduced “click” loudness rating by 50% and scalp sensation by 36%, outperforming standard approaches; and 3) Single sensory experimental modifications alone are not sufficient to significantly reduce vertex N100-P200 or sensory perception.

We show that additional experimental modifications above noise masking alone are needed to reduce the N100-P200 after supratheshold TMS to dlPFC (Figure 3A-G). Compared to prior studies that suggest that noise masking alone can minimize the sensory TEP [2,19], our study differs by intensity and brain target. Regarding intensity, compared to previous work that focused on subthreshold intensities (90% rMT: [2,19]), our suprathreshold stimulation protocol (120% rMT) better mimicked clinical stimulation parameters [64], but is more difficult to mask [18,40,41,65]. In regards to brain target, while other studies have explored sensory suppression after TMS to the primary motor [19] or premotor [2] targets, here we focus on the dlPFC, which may have different sensory contributions to the vertex N100-P200 compared with motor targets [43,66].

While these differences in intensity (80 vs. 120% rMT) or brain target (premotor/M1 vs. dlPFC) may partially explain the inability to fully suppress the vertex N100-P200, other factors may be contributing. The link between sensory potentials, brain target, and intensity is not yet clear [43]. TEPs from M1 stimulation highly correlate with those from non-brain regions (shoulder), regardless of stimulation intensity (120% vs 80% rMT) [18], suggesting that peripherally evoked contributions to the TEP may be considerable regardless of stimulation intensity or target. Multiple studies have demonstrated that N100-P200 components can persist, both for subthreshold and suprathreshold M1 stimulation, even after suppression of TMS click perception [65,67]. Therefore, we cannot conclude that auditory suppression protocols will be effective for all designs, nor that auditory perception of the “click” will be an effective indication of suppression of sensory components in the TEP. We may be yet to find the most effective sensory suppression protocol for all designs and populations. However, our proposed novel combination of sensory reduction procedures (ATTENUATE), which includes extra auditory reduction (with the use of over-the-ear protection), foam, and predictable timing of TMS pulses, is superior at reducing vertex N100-P200 and sensory perception (loudness and scalp feeling) compared to standard experimental procedures.

### 4.1. Effect of a foam spacer on the sensory TEP

We find that a foam spacer attached to the bottom of the coil had no effect on vertex N100-P200, “click” loudness perception, scalp feeling or pain. Although foam could be contributing to the combined effectiveness of ATTENUATE, there was no impact when other modifications were matched (Figure 3H-N). Foam has been suggested to reduce bone conduction of the TMS “click” sound [23] and shown to be effective when used in combination with auditory suppression methods [41]. As such, foam padding between the coil and scalp has become standard procedure to help suppress the vertex N100-P200. However, foam also increases coil to cortex distance and there are no guidelines for adjusting stimulation intensity to account for this increased distance or for reporting whether foam was used in determining motor thresholds. Coil to cortex distance has a strong influence on induced electric field in cortex [68] – enough to significantly increase MT determination [41], as also observed in the current study (Table S1). The lack of reduction in vertex N100-P200 with foam in our conditions when other factors were matched could be due to this increased intensity of TMS with compared to without foam. Interestingly, we also observed no difference in early localized TEP, suggesting that the adjusted rMT with foam likely resulted in a matched induced electrical field, potentially diminishing the argument that the adjusted rMT accounts for our lack of suppression. Overall, our results suggest that foam may not be alone effective for reducing sensory confounds. If used it is important that stimulation intensity is adjusted to account for the increased distance from coil to cortex. Furthermore, this adjusted rMT should be reported in future studies to allow further analysis into this critical question.

### 4.2. Non-modal or multimodal component contributions to the TEP

The vertex N100-P200 complex has been described as an auditory component (see AEP; [23,44]), but it is likely to have multimodal sensory contributions. Although observed in the TEP, vertex N100-P200 complexes with similar/matching time course of peak latencies and similar source activations have been more rigorously examined and described in response to sensory stimuli other than the TMS “click” sound. Many of these studies describe multisensory or cross-modal impacts on the vertex N100-P200 [69–71], suggesting that it is not modality specific and instead largely determined by the intrinsic saliency of the stimulus and its task relevance [28,72].

In TMS-EEG experiments, it is difficult to distinguish between unimodal auditory and non-modal or multimodal sensory contributions to the vertex N100-P200. In addition, it is unclear if sensory contributions are likely to sum linearly. In light of this, sensory suppression protocols for TMS-EEG may be more effective if the vertex N100-P200 is assumed to be multimodal. Our proposed ATTENUATE procedure may demonstrate additional benefit over a *Standard masking* procedure due to over-the-ear auditory masking or saliency reduction through predictable ISI of the TMS pulses. Of note is that neither *auditory masking* (even with over-the-ear protection) nor *predictable ISI* timing was more than borderline or minimally effective at suppressing vertex N100-P200 when used alone. Instead, maximal suppression was achieved when combining *auditory masking* and *predictable ISI* timing, suggesting that a combined sensory masking and sensory attenuation protocol is most effective.

### 4.3. Is sensory suppression the most effective strategy for reducing vertex N100-P200

One clear limitation of our results is that neither perceptual ratings nor vertex N100-P200 were fully eliminated. It should be noted that our design was intended to evoke a large vertex N100-P200 by using suprathreshold stimulation (120% rMT) and with a stimulation target that is known to induce significant sensory artifact [66]. Future work should examine the efficacy of the ATTENUATE protocol across stimulation intensities and targets. ATTENUATE may fully eliminate vertex N100-P200 at lower stimulation intensities or other stimulation targets, but this is outside the scope of the current work and will need to be investigated experimentally.

Furthermore, when the study design allows, sensory suppression techniques should be considered only after other experimental options such as active controls. For instance, if the experimental question allows for conditions with matched intensity, matched sensory suppressive protocols, and target locations with active TMS, then perception and cortical sensory components in the TEP should also be matched. Although this design is optimal, it is not feasible for many studies, either due to time or other constraints. Alternatively, one can isolate the sensory contributions to the TEP using sensory-matched sham protocols. Although it is difficult to match the sensory experience of active TMS with sham TMS, the topography and time course of evoked sensory potentials may be similar [18] enough to employ an ICA-based technique for removal [40,73]. Indeed, a combination of sensory suppression, such as ATTENUATE, coupled with sensory-matched sham TMS may be most effective for reducing the impact of sensory confounds while ensuring that residual sensory contributions to TEP can be more easily identified.

### 4.4. Sensory potentials and pain perception

Our results suggest that perception of pain due to TMS may be relevant to the vertex N100-P200 complex. This result is perhaps unsurprising as previous work has demonstrated substantial overlap between auditory/somatosensory responses and activity in a ‘pain matrix’ network with nociceptive stimuli applied to the skin [28]. Due to a high correlation between the response to sensory and nociceptive stimulation as well as sensory/nociceptive responses and saliency ratings, the authors suggested that sensory responses and pain matrix activity may be best characterized as stimulus saliency-related network activity. Although our correlation and regression analyses were exploratory, this work suggests that reducing the saliency of TMS, including minimization of pain, should be investigated to minimize the vertex N100-P200.

## 5. Future directions

While this work provides critical improvements in sensory suppression during TMS studies, several important questions remain. Given the wide variety of acoustic and somatosensory responses to TMS, the ATTENUATE protocol should be tested with a range of stimulation intensities, coils, and brain targets to establish its efficacy for different stimulation environments. It is also important to explore how ATTENUATE performs in patients with sensory deficits such as hearing loss and sensory processing disorders. Additionally, our data suggest that predictability of TMS pulse timing can contribute to amplitude suppression in sensory TEP, building on prior work showing MEP attenuation with predictable M1 stimulation [45,46]. Although we did not observe a cumulative effect on the TEP using 80 unjittered single pulses of TMS, the sensory predictive suppressive effect should be examined with more single pulses and with a range of unjittered ISIs to ensure that the unjittered protocol does not induce a buildup of brain changes (i.e. neuroplasticity). Sensorimotor prediction for the timing of sensory events is well documented [See [74] and [75] for reviews], and may account for motor and sensory attenuation with predictable TMS pulse timing. However, the suggestion that the principles of sensorimotor timing can be used to optimize non-motor and non-sensory TEP is novel to the best of our knowledge. Future work should compare the effects on the TEP of task-relevant sensorimotor experience [46,83], readiness-to-act [84,85], and interval and phase timing in rhythmically predictable TMS sequences [74,76–82], as variables in the TMS sensory predictive suppressive effect.

## 6. Conclusions

We investigated the electrophysiological and perceptual consequences of applying different sensory suppression protocols with suprathreshold TMS to dlPFC. We find that ATTENUATE outperforms the *Standard masking* protocol for reducing both the vertex N100-P200 and sensory perception. Further, our data support that *auditory suppression, foam spacing*, or *pulse timing* alone are not sufficient to reduce the vertex N100-P200, likely due to the non-modal or multimodal contributions of the sensory experience of the TMS pulse.

## Acknowledgements

We extend gratitude to all of our research participants. We would also like to acknowledge the generous contributions of the members of the Personalized Neurotherapeutics Laboratory for helpful feedback on the manuscript and throughout the course of the study.

This research was supported by the National Institute of Mental Health under award number R01MH126639 and a Burroughs Wellcome Fund Career Award for Medical Scientists (CJK).

JMR was supported by the Department of Veterans Affairs Office of Academic Affiliations Advanced Fellowship Program in Mental Illness Research and Treatment, the Medical Research Service of the Veterans Affairs Palo Alto Health Care System and the Department of Veterans Affairs Sierra-Pacific Data Science Fellowship.

## Appendix A

Supplementary data related to this article can be found online.

## Data availability

Data will be available on NIMH Data Archive within 6 months of publication.

