## Supplementary for "Experimental Suppression of TMS-EEG Sensory Potentials"

**Supplementary materials for:**
Ross JM, Sarkar M, Keller CJ. Experimental Suppression of TMS-EEG Sensory Potentials. [Journal] [In prep]

### S1.1 Preprocessing of TEPs

Data for each channel were epoched with respect to the TMS pulse (-500 to 500ms) and baseline corrected by subtracting the mean amplitude pre-stimulus (-500 to -100ms). Continuous data were reviewed visually for channels that would be problematic for ICA, and only these channels were removed. The average number of channels removed was less than 1 channel (0.68±1.8 channels). The most common reason for channel removal was poor/unstable contact with the scalp. To remove the earliest/highest amplitude TMS-induced electrical artifact, data were zero-padded from -14 to 14ms, following our previous work [40]. Epochs were visually reviewed to identify and reject those epochs that were clearly artifactual (average number of epochs deleted was 2.53±2.2 epochs, or 3.20±2.9%). The most common reason for epoch deletion was an eye blink overlapping with TMS pulse.

To separate EEG patterns that may be artifactual, fast independent component analysis (fICA [60]) was performed at two different stages. After each stage of fICA, a semi-automated artifact detection algorithm incorporated into the open source TMS-EEG Signal Analyzer (TESA v1.1.1) extension for EEGLAB [59] (http://nigelrogasch.github.io/TESA/) was used to classify and visually inspect components based on their scalp topography, time course, and power spectrum. In the first stage, only high amplitude TMS pulse artifacts were removed as this signal masks any other underlying brain and non-brain components and can interfere with a clean ICA decomposition [61,62]. Average number of components removed at this stage was <1 (0.93±0.4 components, for a removal of 0-2.59% of the data across all subjects). Next, data were interpolated for previously zero-padded time window around the TMS pulse using linear interpolation, notch filtered (57-63 Hz), band pass filtered (forward-backward 4th order Butterworth filter, 1-100 Hz), and referenced to the global average. Missing/removed channels were then interpolated using spherical interpolation prior to further preprocessing. All experimental conditions were then merged in order to apply a common ICA decomposition across all conditions. To identify and remove remaining non-brain artifacts including eye movement/blink, single electrode noise, TMS evoked muscle [62], and cardiac beats (EKG), a second round of fICA was performed on the merged data. EKG artifacts were only removed if they appeared to be not mixed with brain signal. The average number of remaining brain components following all data cleaning was 15.67±4.9 components. Finally, to minimize the impact of electrical noise and non-brain artifacts, the remaining cleaned data were low pass filtered (4th order Butterworth filter at 50 Hz). Experimental conditions were then unmerged for subsequent analysis.

### S1.2 Quantification of TEPs

To perform analyses across experimental conditions, the time windows for the early TEP (<100), N100, and P200 needed to be determined. To do so, we examined the group averaged global mean field potential (GMFP) in the *No masking* condition, as previously described [63,64]. Twenty subjects had this *No masking* condition (Supplementary tables S2-3 for which conditions were collected for each subject, and for *No masking* condition Figure S1 for individual subject TEPs and Figure S2 for TEPs averaged across subjects). We utilized the 14-300 ms post-TMS window as <14ms represents interpolated data from the stimulation artifact and >300ms does not involve the N100-P200 time window and typically comprises low voltage. To identify relevant TEP peaks in this 14-300ms time window, an automated peakfinder algorithm [65] processed the averaged GMFP time series (N=20) and outputted peaks >2SD above baseline (-400 to -14ms). Four peaks were identified at the following latencies: 50, 122, 175, and 226 ms. The P200 can be seen in these data as having two small peaks at 175 and 226 ms (Figure S3). Local minima between potentials were at 86, 138, and 285 ms (Figure S3). From these group average *No masking* condition GMFP peak/trough times, three time windows were selected for the between-condition analyses. These time windows were selected to capture the entire potentials, with window boundaries defined by troughs in the GMFP: 14-86 ms (early window), 86-138 ms (mid-latency), and 138-285 ms (longer latency).

To identify a region of interest (ROI) for the N100 and P200 time windows, group topoplots from the *No masking* condition (N=20) were utilized. For each time window (86-138ms, 138-285ms), topoplots were averaged over the time window and baseline subtracted (-400 to -100ms). The electrodes demonstrating maximal difference from baseline and shared between the N100 and P200 time windows included the following: FC1, FCz, FC2, C1, Cz, C2 (Figure S4 for individual subject and Figure S5 for group averaged topoplot). To compare vertex N100-P200 across experimental conditions, TEPs were generated as averages over each ROI. Local mean field potential (LMFP) was calculated for the ROI and the area under the curve (AUC) of the LMFP was quantified for the appropriate time windows. Supporting that our time windows and ROI capture the vertex N100-P200 complex, we observed a strong correlation between vertex N100 and P200 (Figure S7A,B; area under the curve (AUC) of LMFP; $r(19)=0.91$, $p=0.00000004$; regression: $F(1,19)=81.68$, $p=0.00000004$; $R^2 = 0.82$).

| Subject | Age | Dominant Hand | Gender | Optimized Coil Angle | rMT without foam | rMT with foam |
|---|---|---|---|---|---|---|
| 1007 | 28 | R | F | 45 | 74 | 89 |
| 1008 | 24 | R | M | 45 | 57 | 61 |
| 1010 | 27 | R | M | 20 | 64 | 73 |
| 1011 | 19 | R | F | 80 | 66 | 83 |
| 1015 | 33 | R | M | 90 | 56 | 60 |
| 1024 | 49 | R | F | 0 | 94 | >100* |
| 1028 | 31 | R | F | 45 | 82 | 86 |
| 1043 | 64 | R | F | 90 | 62 | 68 |
| 1044 | 46 | R | F | 40 | 70 | 84 |
| 1054 | 43 | R | F | 45 | 80 | 87 |
| 1062 | 61 | R | F | 20 | 72 | 79 |
| 1064 | 62 | R | M | 45 | 54 | 57 |
| 1073 | 64 | R | F | 25 | 80 | 81 |
| 1075 | 44 | R | F | 45 | 62 | 68 |
| 1077 | 54 | R | F | 90 | 74 | 82 |
| 1082 | 45 | L | F | 45 | 73 | 79 |
| 1132 | 62 | R | M | 45 | 68 | 74 |
| 1143 | 24 | R | M | 100 | 55 | 63 |
| 1145 | 45 | R | F | 60 | 65 | 72 |
| 1147 | 43 | R | M | 80 | 75 | 89* |
| 1148 | 55 | R | F | 45 | 59 | 61 |

**Table S1.** Subject demographics (age in years, handedness, gender) and stimulation details (coil angle measured in degrees from cap midline, rMT without and with foam in %MSO) for all N=21 subjects. Due to limitations on experiment time, all conditions were not completed by all subjects. For missing conditions, see Table S2-S3. Stimulation intensity for each condition was set to 120% rMT. *Due to high subject rMT, stimulation was administered at 100% MSO instead of 120% rMT.

| **Protocol** | *No masking* | | | | | | | | *Standard masking* | | ATTENUATE | |
|---|---|---|---|---|---|---|---|---|---|---|---|---|
| **Individual mods.** | *No noise* | | *Noise* | | *Noise + over-the-ear protection /No foam* | | *Foam/Jittered* | | | | *Unjittered* | |
| | N100 | P200 | N100 | P200 | N100 | P200 | N100 | P200 | N100 | P200 | N100 | P200 |
| 1007 | 126.68 | 510.06 | 212.07 | 813.93 | 139.50 | 623.54 | | | 155.97 | 534.75 | 106.47 | 553.11 |
| 1008 | 464.80 | 762.32 | 191.04 | 370.09 | 271.13 | 676.60 | 487.94 | 1009.86 | 281.36 | 415.06 | 185.71 | 399.85 |
| 1010 | 114.31 | 586.81 | 194.44 | 282.08 | 227.25 | 375.11 | 146.65 | 457.15 | 206.90 | 366.30 | 126.63 | 194.99 |
| 1011 | | | | | | | 324.84 | 373.61 | | | 288.22 | 232.01 |
| 1015 | 109.65 | 406.95 | 103.11 | 327.22 | 176.93 | 359.29 | 205.07 | 348.09 | 113.52 | 333.64 | 201.58 | 356.67 |
| 1024 | 115.31 | 459.64 | 199.03 | 814.67 | 180.64 | 551.07 | 75.61 | 455.84 | 362.64 | 1141.96 | 108.67 | 267.63 |
| 1028 | 182.62 | 682.11 | 102.38 | 484.23 | 101.44 | 452.08 | 126.58 | 399.52 | 111.28 | 516.75 | 86.46 | 310.53 |
| 1043 | 276.33 | 512.33 | 169.81 | 273.44 | 63.41 | 135.32 | 65.83 | 239.07 | 201.16 | 327.99 | 113.85 | 273.30 |
| 1044 | 200.28 | 740.08 | 158.29 | 503.38 | 180.93 | 726.24 | 128.14 | 331.22 | 244.38 | 688.42 | 158.78 | 489.26 |
| 1054 | 177.76 | 622.51 | 248.31 | 558.98 | 105.77 | 391.49 | 101.78 | 369.95 | 182.51 | 498.48 | 107.02 | 271.04 |
| 1062 | 107.00 | 319.23 | 168.66 | 518.38 | 153.35 | 414.04 | 111.26 | 277.33 | 167.05 | 459.92 | 125.74 | 232.37 |
| 1064 | 152.69 | 485.22 | 61.95 | 234.13 | 45.26 | 124.26 | 52.07 | 161.12 | 62.66 | 212.08 | 81.02 | 164.66 |
| 1073 | 252.55 | 647.29 | 75.90 | 321.01 | 123.09 | 243.74 | 114.40 | 290.46 | 127.11 | 300.99 | 197.86 | 162.64 |
| 1075 | 226.05 | 542.05 | 223.45 | 319.83 | 144.13 | 221.37 | 109.91 | 236.65 | 191.10 | 368.09 | 124.19 | 174.30 |
| 1077 | 237.24 | 614.87 | 109.89 | 416.48 | 233.38 | 559.91 | 51.94 | 256.14 | 164.64 | 548.74 | 131.48 | 219.19 |
| 1082 | 110.27 | 721.84 | 118.23 | 791.51 | 92.89 | 590.23 | 123.33 | 613.29 | 155.85 | 812.78 | 92.53 | 251.88 |
| 1132 | 376.09 | 901.91 | 318.35 | 645.20 | 270.18 | 556.12 | 266.51 | 677.67 | 322.18 | 806.17 | 95.70 | 294.23 |
| 1143 | 780.33 | 1375.92 | 394.50 | 678.66 | 536.48 | 810.10 | 544.95 | 832.77 | 374.78 | 461.21 | 297.11 | 473.50 |
| 1145 | 78.10 | 498.88 | 74.08 | 262.10 | 92.03 | 288.93 | 82.75 | 253.95 | 64.40 | 275.41 | 104.02 | 253.62 |
| 1147 | 1076.92 | 2088.35 | 893.09 | 1570.72 | 462.73 | 883.08 | 479.40 | 1069.45 | 797.60 | 1460.39 | 209.64 | 634.31 |
| 1148 | 920.53 | 1075.30 | 468.97 | 628.21 | 380.51 | 440.04 | 544.93 | 780.30 | 440.59 | 748.89 | 119.86 | 342.41 |

**Table S2.** Vertex N100-P200 LMFP AUC for individual subjects (trial average), for all three sensory suppression protocols (N=20) and all individual modifications (3 auditory conditions (N=20), 2 foam conditions (N=20), 2 jitter conditions (N=20)). Subject 1011 was only included in *Jittered/Unjittered* contrast due to missing conditions. Subject 1007 was not included in *No foam/Foam* contrast or *Jittered/Unjittered* contrast due to missing conditions.

| Protocol | *No masking* | | | | | | | | | | | | *Standard masking* | | | ATTENUATE | | |
|---|---|---|---|---|---|---|---|---|---|---|---|---|---|---|---|---|---|---|
| Individ. mods. | *No noise* | | | *Noise* | | | *Noise + over-the-ear protection /No foam* | | | *Foam/Jittered* | | | | | | *Unjittered* | | |
| | Loudness | Scalp | Pain | Loudness | Scalp | Pain | Loudness | Scalp | Pain | Loudness | Scalp | Pain | Loudness | Scalp | Pain | Loudness | Scalp | Pain |
| 1007 | 5 | 5 | 5 | 3 | 6 | 6 | 1.5 | 6 | 6 | | | | 2 | 6 | 5 | 3 | 6 | 6 |
| 1008 | 7 | 8 | 1 | 0 | 6 | 1 | 0 | 3 | 0 | 0 | 7 | 1 | 1 | 3 | 0 | 0 | 5 | 0 |
| 1010 | 3 | 8 | 4 | 2 | 4 | 1 | 1 | 7 | 4 | 0 | 2 | 0 | 2 | 7 | 4 | 1 | 1 | 0 |
| 1011 | | | | | | | | | | 3 | 4 | 4 | | | | 5 | 6 | 6 |
| 1015 | 5 | 7 | 2 | 2 | 4 | 1 | 2 | 6 | 2 | 0 | 5 | 2 | 4 | 8 | 4 | 3 | 7 | 5 |
| 1024 | 6 | 6 | 1 | 5 | 2 | 1 | 5 | 4 | 3 | 3 | 3 | 1 | 6 | 1 | 1 | 3 | 1 | 1 |
| 1028 | 5 | 8 | 0 | 4 | 6 | 0 | 3 | 6 | 0 | 3 | 6 | 0 | 5 | 7 | 0 | 3 | 7 | 0 |
| 1043 | 1 | 2 | 1 | 1 | 1 | 1 | 0.5 | 0.5 | 1 | 1 | 0.5 | 3 | 1 | 2 | 1 | 1 | 1 | 1 |
| 1044 | 3 | 3 | 1 | 2 | 4 | 4 | 3 | 5 | 3 | 1 | 1 | 3 | 3 | 5 | 1 | 3 | 5 | 3 |
| 1054 | 3 | 5 | 1 | 3 | 4 | 1 | 3 | 6 | 3 | 3 | 2 | 1 | 5 | 4 | 2 | 3 | 4 | 3 |
| 1062 | 7 | 4 | 3 | 5 | 5 | 3 | 6 | 5 | 3 | 4 | 5 | 2 | 5 | 5 | 3 | 4 | 5 | 3 |
| 1064 | 1 | 1 | 0 | 0.5 | 0.5 | 0 | 1 | 0.5 | 0 | 0.5 | 1 | 0.5 | 1 | 1 | 0 | 0 | 0.25 | 1 |
| 1073 | 3 | 9 | 1 | 5 | 5 | 4 | 3 | 4 | 1 | 2 | 4 | 1 | 3 | 4 | 0 | 0 | 2 | 0 |
| 1075 | 2 | 8 | 2 | 2 | 3 | 6 | 0 | 3 | 3 | 0 | 2 | 1 | 1 | 3 | 2 | 0 | 5 | 0 |
| 1077 | 4 | 10 | 1 | 2 | 9 | 3 | 3 | 10 | 2 | 1 | 6 | 2 | 2 | 6.5 | 0 | 1 | 6 | 3 |
| 1082 | 3.5 | 6 | 0 | 3.5 | 6 | 0 | 1 | 4.5 | 0 | 4 | 5 | 0 | 3.5 | 5 | 0 | 1 | 4 | 0 |
| 1132 | 3 | 7 | 4 | 6 | 9 | 6 | 4 | 5 | 4 | 2 | 7 | 4 | 3 | 6 | 1 | 3 | 6 | 0 |
| 1143 | 5 | 6 | 2 | 4 | 5 | 2 | 3 | 4 | 1 | 3 | 4 | 2 | 3 | 6 | 5 | 4 | 6 | 6 |
| 1145 | 5 | 7 | 2 | 2 | 7 | 5 | 1 | 2 | 2 | 2 | 2 | 2 | 4 | 7 | 3 | 3 | 3 | 4 |
| 1147 | 5 | 8 | 7 | 3 | 7 | 7 | 2 | 7 | 7 | 3 | 2 | 2 | 4 | 6 | 3 | 3 | 4 | 4 |
| 1148 | 8 | 8 | 0 | 7 | 7 | 0 | 2 | 2 | 0 | 4 | 4 | 0 | 3 | 3 | 6 | 3 | 3 | 2 |

**Table S3.** Perceptual ratings for individual subjects for all three sensory suppression protocols (N=20) and all individual modifications (3 auditory conditions (N=20), 2 foam conditions (N=20), 2 jitter conditions(N=20)). Rating are perception of TMS “click” loudness, scalp sensation, and pain, on 0-10 ratings scales. Subject 1011 was only included in *Jittered/Unjittered* contrast due to missing conditions. Subject 1007 was not included in *No foam/Foam* contrast or *Jittered/Unjittered* contrast due to missing conditions.

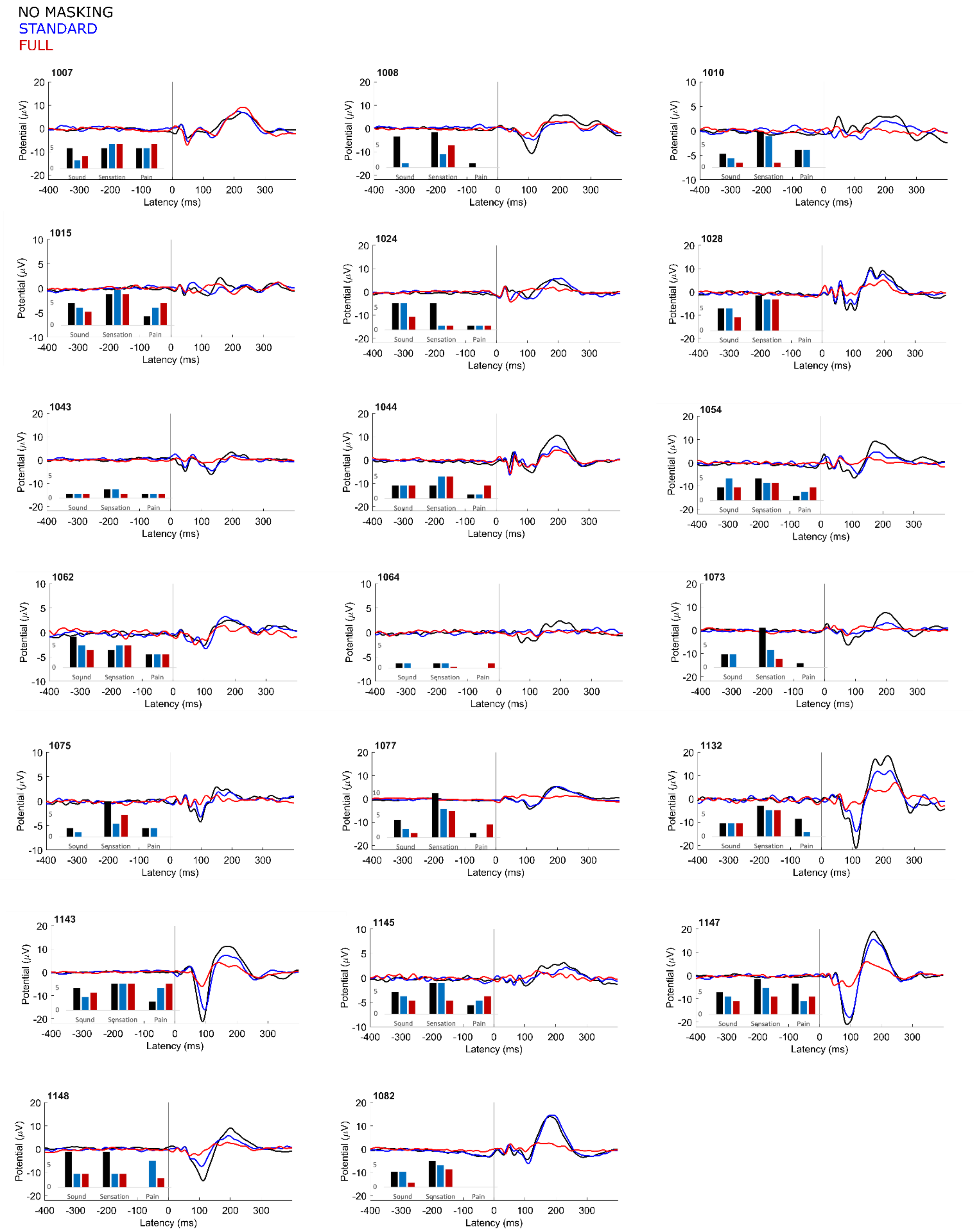


**Figure S1.** Individual subject TEPs and perceptual ratings. TEPs were averaged across the vertex ROI (C1, Cz, C2, FC1, FCz, FC2) for *No masking*, *Standard masking*, and ATTENUATE conditions. Perceptual ratings were given for 0-10 scales of loudness of TMS "click," intensity of scalp sensation, and pain.

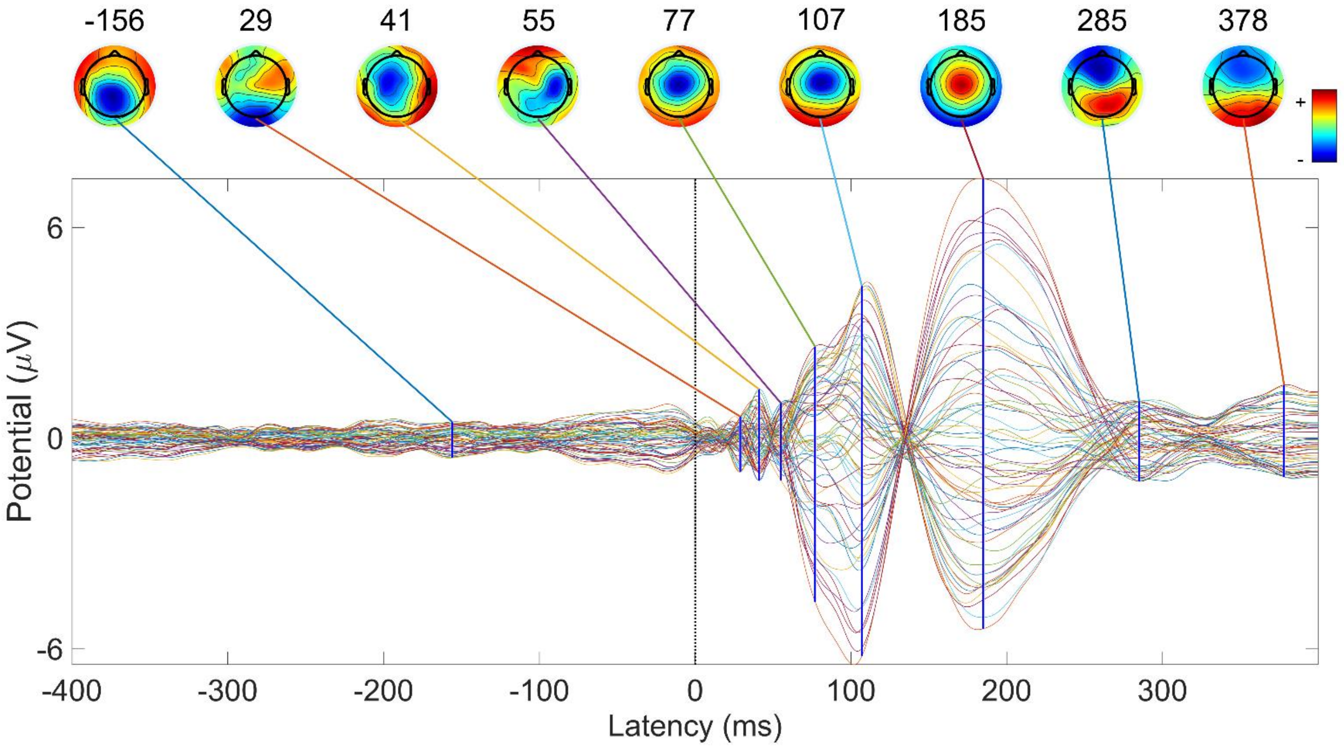


**Figure S2.** TEP averaged across all subjects with the *No masking* condition (N=20).

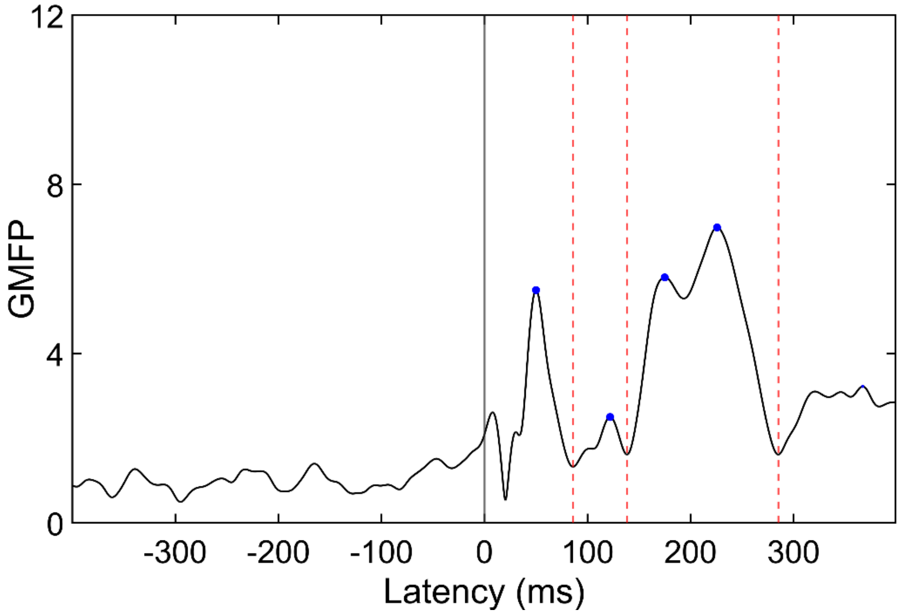


**Figure S3.** GMFP timeseries averaged across all subjects with *No masking* condition (N=20). Peaks in the GMFP that were greater than 2 standard deviations above baseline are identified in blue, with local minima between peaks marked in red.  Local minima between potentials were at 86, 138, and 285 ms. From these group average peak/trough times, three time windows were selected for all between condition analyses, with window boundaries defined by troughs: 14-86 ms (early window), 86-138 ms (mid-latency), and 138-285 ms (longer latency).

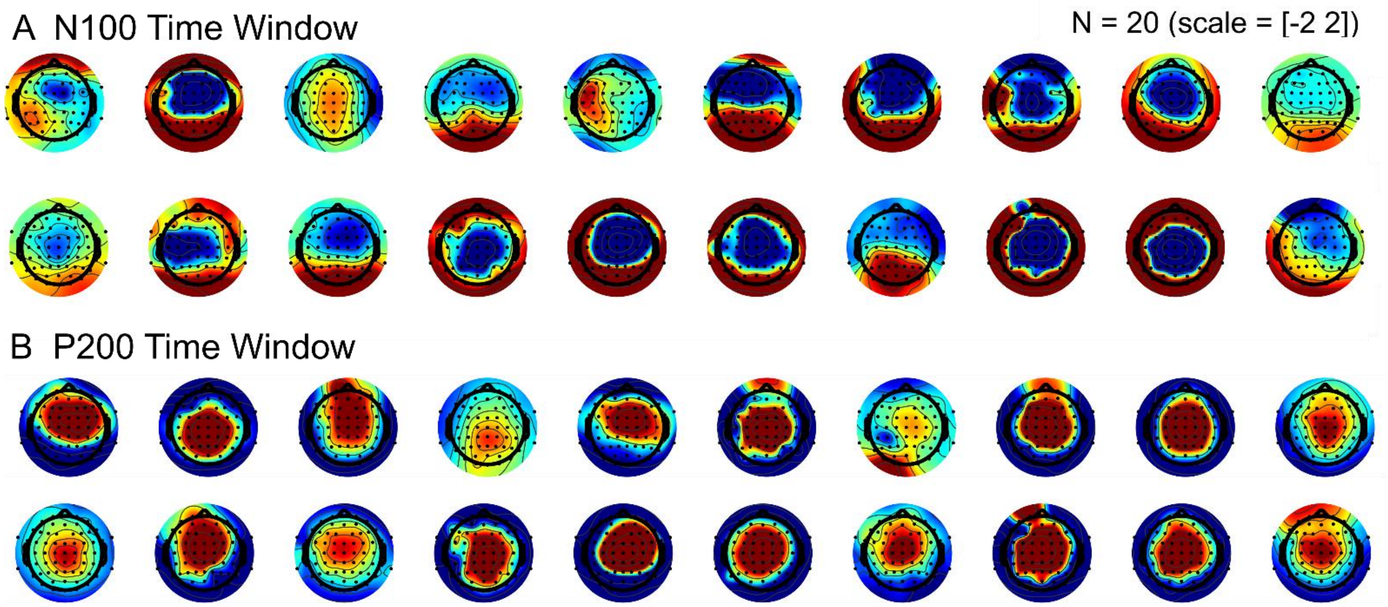


**Figure S4.** Topoplots of all individual subjects with *No masking* condition in N100 and P200 time windows, defined using peak/trough analysis (Figure S3). For each time window (86-138ms, 138-285ms), topoplots were averaged over the time window and baseline subtracted (-400 to -100ms).

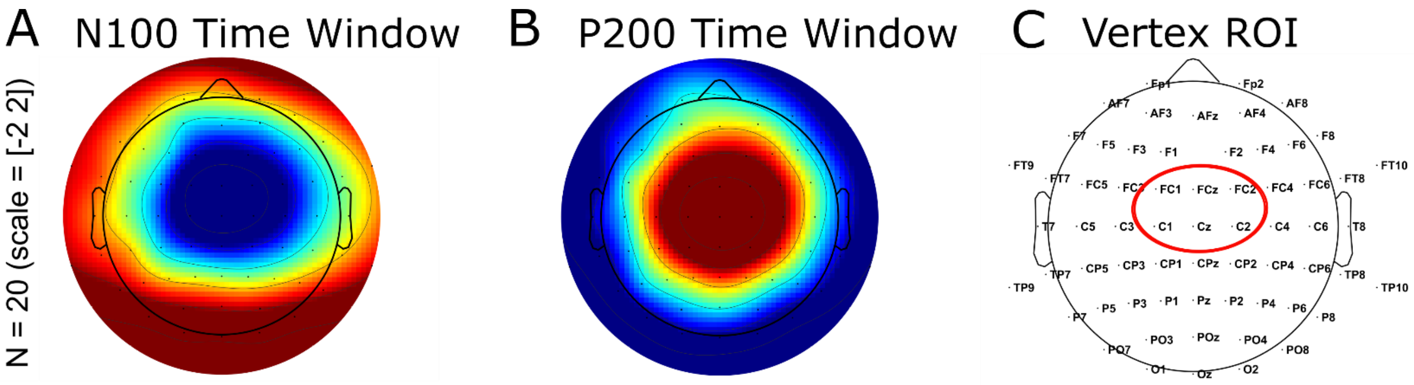


**Figure S5.** (A-B) Topoplots averaged over all subjects with *No masking* condition (N=20) in (A) N100 and (B) P200 time windows, defined using peak/trough analysis (Figure S3). (C) The electrodes demonstrating maximal difference from baseline and shared between the N100 and P200 time windows included the following: FC1, FCz, FC2, C1, Cz, C2. To compare vertex N100-P200 across experimental conditions, TEPs were generated as averages over this vertex ROI. LMFP was calculated for the ROI/time windows relevant to each experimental contrast, and LMFP area under the curve (AUC) was calculated.

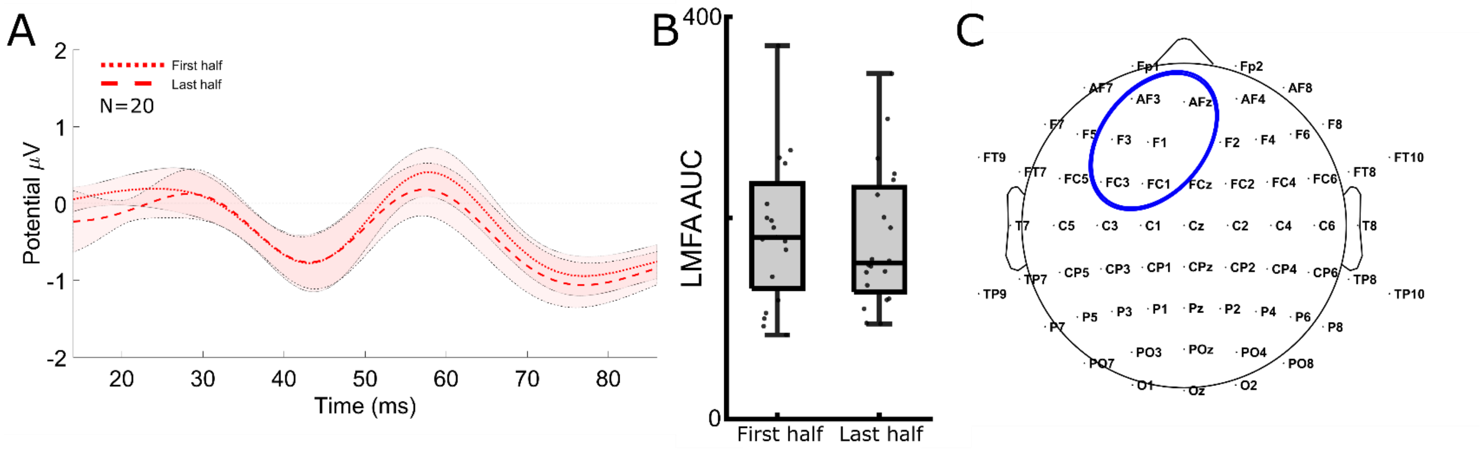


**Figure S6.** We find no cumulative effect on excitability in early (14-86 ms) local (AF3, AFz, F3, F1, FC3, FC1) TEP with the *Unjittered* condition. (A) Early local TEP from first 40 trials and last 40 trials of the *Unjittered* condition (averaged across all subjects with this condition, N=20). (B) Using the *Unjittered* protocol with 2 second ISI resulted in no difference in LMFP between the first 40 and last 40 trials (T=1.2542, DF=19, p=0.2250, CI=-11.4509, 45.6910). (C) ROI local to dlPFC stimulation target used in all local analyses: AF3, AFz, F3, F1, FC3, FC1.

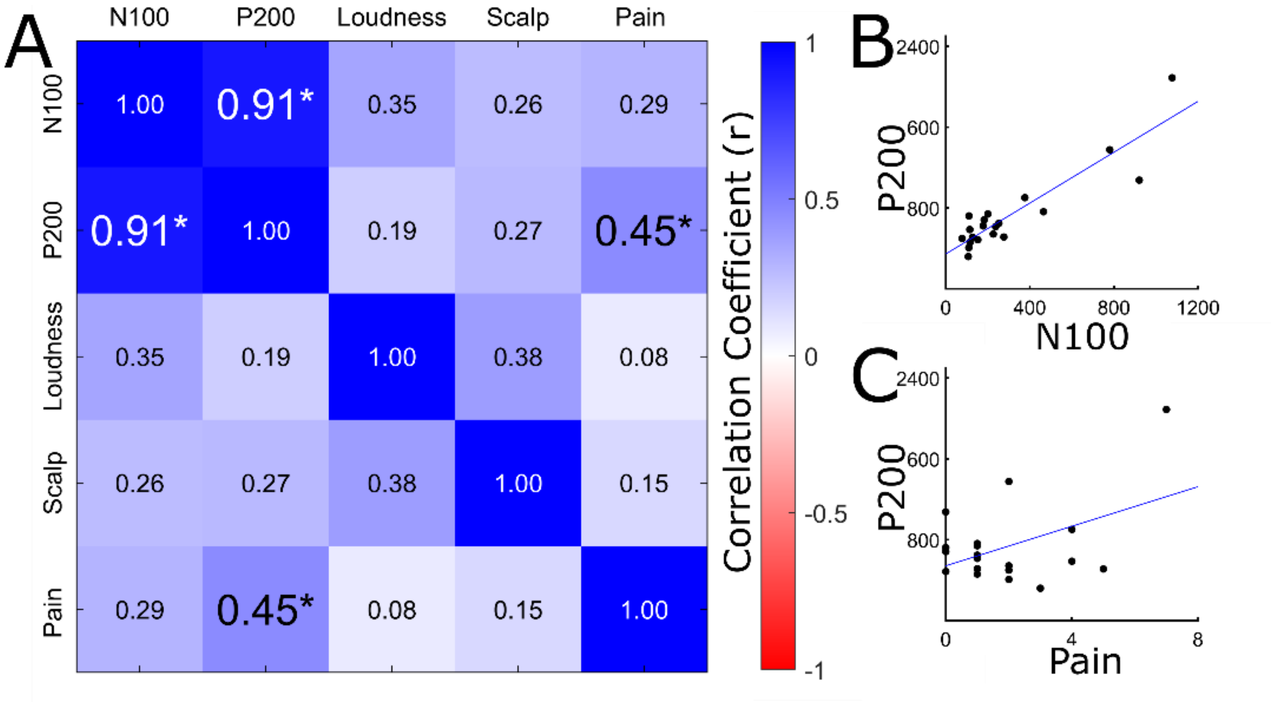


**Figure S7: Pain may be contributing to vertex N100-P200.** (A) Pearson correlation matrix of perceptual ratings and vertex N100-P200 LMFP AUC. We find a relationship between vertex N100 and vertex P200, and between pain rating and vertex P200. (B-C) Regression analysis between vertex N100 and vertex P200 LMFP AUC (B) and between perception of pain and vertex P200 LMFP AUC (C). We find that pain may be contributing to the vertex P200.
